# Reorientation and propulsion in fast-starting zebrafish larvae: an inverse dynamics analysis

**DOI:** 10.1101/570812

**Authors:** Cees J. Voesenek, Remco P. M. Pieters, Florian T. Muijres, Johan L. van Leeuwen

## Abstract

Most fish species use fast starts to escape from predators. Zebrafish larvae perform effective fast starts immediately after hatching. They use a C-start, where the body curls into a C-shape, and then unfolds to accelerate. These escape responses need to fulfil a number of functional demands, under the constraints of the fluid environment and the larva’s body shape. Primarily, the larvae need to generate sufficient escape speed in a wide range of possible directions, in a short-enough time. In this study, we examined how the larvae meet these demands. We filmed fast starts of zebrafish larvae with a unique five-camera setup with high spatiotemporal resolution. From these videos, we reconstructed the three-dimensional swimming motion with an automated method and from these data calculated resultant hydrodynamic forces and, for the first time, 3D torques. We show that zebrafish larvae reorient mostly in the first stage of the start by producing a strong yaw torque, often without using the pectoral fins. This reorientation is expressed as the body angle, a measure that represents the rotation of the complete body, rather than the commonly used head angle. The fish accelerates its centre of mass mostly in stage 2 by generating a considerable force peak while the fish “unfolds”. The escape direction of the fish correlates strongly with the amount of body curvature in stage 1, while the escape speed correlates strongly with the duration of the start. This may allow the fish to independently control the direction and speed of the escape.

**Summary statement:** Fish larvae can independently adjust the direction and speed of their fast start escape response, a manoeuvre crucial for survival.

## Introduction

The fast start is an important manoeuvre in the motion repertoire of many fish species across developmental stages (Domenici and Blake, 1997; Hale et al., 2002). Fast starts are commonly divided into two types by the shape changes of the fish during the motion: the S-start and the C-start. This article concerns the C-start, which is mainly used to escape from (potential) threats (Walker et al., 2005), and in some species for prey capture (Wöhl and Schuster, 2007). It involves the fish bending itself into a C-shape, and then unfolding to produce a strong acceleration and a change of direction (Hertel, 1966; Weihs, 1973). This motion is often considered to consist of three stages (Domenici and Blake, 1997; Hertel, 1966; Weihs, 1973): stage 1, where the fish bends into a C-shape; stage 2, where the fish unfolds; and stage 3, the remainder of the motion – continuous swimming or coasting. In this study, we look at the first two stages of the C-start – we do not consider the highly variable third stage.

For the fast start to contribute to the survival of the larvae, the stages need to satisfy a number of functional demands (Voesenek et al., 2018). The primary demand on a start is to escape from a predator (Domenici and Blake, 1997). This requires strong accelerations to create sufficient distance in a short time between the predator and the larvae (Walker et al., 2005). In addition, it requires control over the escape angle, as the relative heading with respect to the predator often determines escape success (Domenici et al., 2011). Since predators may approach from all sides, it is necessary that the larvae can produce a large range of possible escape directions, both horizontally and vertically. Finally, the threat should be detected early, and the response needs to be well-timed for the escape to be effective (Stewart et al., 2013).

These functional demands should be fulfilled within physical constraints on the body of the larva and the hydrodynamics. Fish larvae need to be able to escape immediately after hatching (Voesenek et al., 2018), while their muscles (Van Raamsdonk et al., 1978), sensory system, and motor control (Fetcho and McLean, 2010) are not fully developed – even within these limits, the larvae need to respond appropriately, quickly, and produce effective motion. Furthermore, to perform effective propulsion as an undulatory swimmer, the larva needs to prepare its body for a propulsive tail-beat by bending into a C (Foreman and Eaton, 1993). To produce thrust, the fish also needs to “prepare” the surrounding water by generating (precursors to) vortices and jets that will contribute to the hydrodynamic forces in stage 2 (Ahlborn et al., 1991; Tytell and Lauder, 2008). In addition, stage 1 prepares the axial muscle for maximum power production by active lengthening of the contralateral side during bending (James and Johnston, 1998).

To meet the functional demands of the fast start, the fish larvae must generate hydrodynamic forces and torques, producing linear and angular accelerations. Different methods have been used to quantify these forces and torques. The motion of the fish and the flow can be quantified with high-speed video images and particle image velocimetry, allowing estimation of momentum changes of the fish and flow (Tytell and Lauder, 2008), or estimation of forces via a reconstructed pressure (Lucas et al., 2017). The reconstructed motion can also be used as input to a computational fluid dynamics method to estimate the forces (Borazjani et al., 2012). Alternatively, the net forces and torques can be reconstructed from kinematics without requiring flow visualisation or fluid-dynamic models, based on inverse dynamics (Van Leeuwen et al., 2015; Voesenek et al., 2016). Since the hydrodynamics are the only source of external forces and torques acting on the fish, we can use the net accelerations of the fish – both linear and angular – to calculate the hydrodynamic forces and torques directly from the kinematics.

The kinematics of the fast start have been characterised in many species (Domenici and Blake, 1993; Fleuren et al., 2018; Kasapi et al., 1993; Müller and Van Leeuwen, 2004). Fast starts have been stated to occur mostly in the horizontal plane (Domenici and Blake, 1997), and most studies investigate two-dimensional kinematics from single-camera high-speed video (e.g. Domenici and Blake, 1993; Harper and Blake, 1990; Hertel, 1966). However, three-dimensional kinematics studies show a vertical motion component in adults (Butail and Paley, 2012; Fleuren et al., 2018; Kasapi et al., 1993) and larval fish (Nair et al., 2015; Stewart et al., 2014). This vertical component is ecologically relevant, since it may influence the effectiveness of predator evasion with the escape response (Stewart et al., 2014).

In this article, we analyse fast starts of zebrafish larvae at 5 days after fertilisation. We filmed fast-start behaviour with a synchronised five-camera setup with high spatial and temporal resolution (Fig. 1A). From these videos, we reconstructed the kinematics in 3D (Fig. 1B,C) and used these data to calculate resultant hydrodynamic forces and torques. Based on the three-dimensional dynamics, we examined how zebrafish larvae meet the functional demands on the fast start. We show that zebrafish larvae produce torques in stage 1 that provide most of the reorientation of the body, while limited propulsion is produced. This is followed by a peak in propulsive force in stage 2, resulting in a strong acceleration of the centre of mass. The turn angle of a start is mostly determined by the amount of body curvature, while the speed at the end of stage 2 is mostly determined by the duration of the start. This allows early-development larvae to perform appropriate escape responses for threats approaching from different directions and at different speeds.

**Fig. 1.**
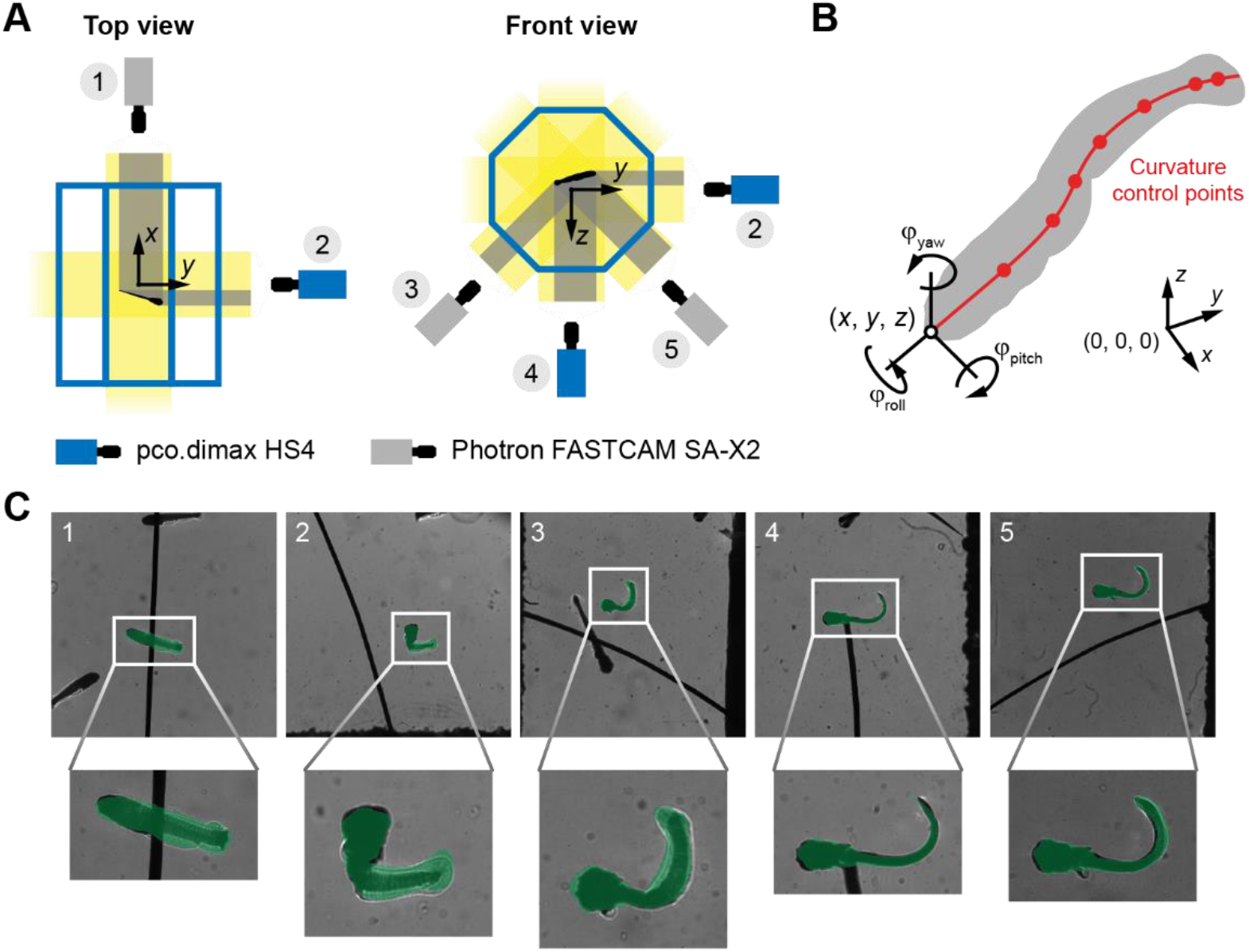
Multi-camera setup and automated tracking method. (A) Sketch of the five-camera setup, top view (left) and front view (right). (B) Parameterised fish model used in the automated tracking method. The model parameters are the 3D position of the snout, the 3D orientation of the head as expressed by the Tait-Bryan angles (roll, pitch, and yaw), and a series of control points for the curvature along the body. Adapted from Voesenek et al. (2016) (C) Overlap between the high-speed video images (grayscale background) and projections of the body model (transparent green). The numbers in the top left corner correspond to the camera numbers in (A).

## Materials and methods

### Animals

We used two batches (from different parents) of 50 wild type zebrafish larvae (*Danio rerio*, Hamilton 1822), bred at the Carus animal facilities of Wageningen University. All fast start sequences were filmed for fish of 5 days post fertilisation, with a body length of 4.2±0.14 mm. We housed each batch in a separate tank, kept at a constant temperature of 27°C. The experimental aquarium was also maintained at 27°C by heating the experimental room. We placed 50 larvae at the same time in the aquarium. The fish were stimulated to perform fast start manoeuvres by approaching them with a horse hair. Sequences where the hair touched the fish were eliminated from analysis, because the resultant forces would not only be from the hydrodynamics, but also from the hair. The influence of the flow induced by the hair is limited: the centre of mass of the larvae hardly moves before the initiation of the start. All experiments were approved by the Wageningen University animal ethics committee.

### Experimental setup

The swimming of larval zebrafish was recorded with a synchronised high-speed video setup with five cameras with different orientations (Fig. 1A). Zebrafish larvae were placed in a glass aquarium in the shape of an octagonal prism (12 mm sides). To limit refraction effects, the cameras were placed perpendicular to the glass from five angles. From the bottom and the right side, we used pco.dimax HS4 cameras (PCO AG, Kelheim, Germany; 2000×2000 pixels). From the back, bottom left, and bottom right side, we used Photron FASTCAM SA-X2 cameras (Photron, Tokyo, Japan; 1024×1024 pixels). All cameras were equipped with 105 mm f/2.8 macro lenses (105 mm f/2.8 FX AF MICRO-NIKKOR and AF-S 105 mm f/2.8G VR Micro, Nikon, Tokyo, Japan) with +5 diopter close-up lenses (DHG Achromat Macro 200(+5), Marumi, Nagano, Japan), mounted on 27.5 mm extension tubes (PK-13, Nikon, Tokyo, Japan). All cameras were recording at 2200 frames per second, synchronised with a pulse generator (9618+, Quantum Composers, Bozeman, Massachusetts, USA). By using a collimated light setup, we created high-contrast shadow images with large depth of field. Collimated light was produced by shining an LED light source (MNWHL4/MWWHL4, Thorlabs Inc., Newton, New Jersey, USA) placed in the focus of a 250 mm lens (250D, Canon, Tokyo, Japan). The light setup was aligned such that the collimated light was parallel with the optical axis of the camera. Since the fish larvae were in an aquarium between the light source and the camera, they projected deep shadows on a brightly lit background image at short shutter speeds (≈10 μs).

### Camera calibration and modelling

We generated calibration points visible in all cameras by moving a sharp-tipped needle through the measurement volume with a computer-controlled micromanipulator (MCL-3, LANG GmbH & Co. KG, Hüttenberg, Germany). The needle was moved through a cuboid volume, at 5×5×5 uniformly spaced points along each dimension. This resulted in 125 images per camera with a known position of the needle tip. In each of these images, we indicated the needle tip manually with a custom Python 3 program.

Camera projections were modelled by a simple affine transform, where we ignored perspective effects. For our camera setup, this is a valid assumption, as the shadows projected onto the sensor by the fish are (theoretically) independent of the distance from the sensor, owing to the collimated light. The affine transform for each camera was parameterised by a 3D translation and the orthonormal basis of the image plane coordinate system (i.e. one outward and two in-plane vectors). From an initial estimate of the camera parameters, we started a constrained optimisation procedure in MATLAB (interior-point algorithm as implemented in `fmincon`; R2016a, The Mathworks, Natick, Massachusetts, USA). Using this procedure, we minimised the sum of squared differences between the clicked image coordinates and the reprojected image coordinates, while maintaining orthonormality (i.e. all vectors perpendicular and of unit length) of the image plane basis vectors.

### Motion reconstruction

The motion of the larvae was reconstructed from the synchronised high-speed video with the method described in Voesenek et al. (2016); it was originally developed in MATLAB, but converted to Python 3. We will briefly summarise the method here, but refer the reader to the original article for more details.

The method is based on a virtual representation of the camera setup and the fish larva. The virtual camera setup was created from the results of the calibration procedure described above. It transforms a point in world coordinates to image plane coordinates for each camera. The fish was represented by a three-dimensional surface model. The shape, position, and orientation of this model were determined by 14 parameters (3 for position, 3 for orientation, 8 for body curvature control points; Fig. 1B) – we ignore dorsoventral curvature, deformation of the median fin fold, and motion of the pectoral fins. For every point in time, we applied the Nelder-Mead optimisation algorithm to these parameters to minimise the difference between virtual images, for which the 3D model was projected onto the virtual cameras, and the real high-speed video images, from which the fish was segmented. The result was a time series of body curvature along the body, position, and orientation that described a three-dimensional surface with optimal overlap (Fig. 1C). We smoothed each of these time series with regularised least squares (Eilers, 2003; Stickel, 2010), with derivatives of order 4, and a smoothing parameter of 100.

The reconstructed time series of parameters uniquely described the 3D shape of the fish. Under the assumption of a constant density across the fish, the mass distribution is known at every point in time. This allowed us to calculate its linear and angular momentum, and therefore the resultant fluid-dynamic forces and torques (Voesenek et al., 2016). In addition, for each frame in each tracked sequence, we determined visually from the bottom camera whether the pectoral fins were abducted or adducted.

### Body angle calculation

We calculated the body angle by integrating angular velocity obtained from the angular momentum. We calculated the angular velocity as **ω** = **I**^−1^**L**, where **ω** is the angular velocity vector in rad s^−1^, I is the moment of inertia tensor, and **L** is the angular momentum vector. We integrated this angular velocity vector with the midpoint rule (Simo and Wong, 1991; Zupan and Saje, 2011) to obtain rotation matrices, with the rotation matrix of the head at the beginning of the start as the initial condition. Finally, we reconstructed the body roll, pitch, and yaw Tait-Bryan angles from these rotation matrices.

### Statistics

For all statistical tests, we used a significance threshold of 0.05. We performed all statistics with MATLAB (R2018b, The Mathworks, Natick, Massachusetts, USA) and the associated Statistics and Machine Learning Toolbox (R2018b, The Mathworks, Natick, Massachusetts, USA). We verified normality of the data with a Kolomogorov-Smirnov test (MATLAB’s `kstest`). To calculate correlation coefficients, we fitted linear models (MATLAB’s `fitlm`). We standardised all data before fitting the model by subtracting its mean and dividing by its standard deviation, which allowed us to use the fit coefficients as correlation coefficients (Schielzeth, 2010). To calculate confidence intervals of the correlation coefficients, we used bootstrapping with 10,000 repetitions, then calculated the 2.5^th^ and 97.5^th^ percentile. The correlation coefficients and their confidence intervals were converted back into slopes by multiplying with *σ_x_/σ_y_*, the ratio of standard deviations.

For the models of the turn angle and speed as a function of the head-to-tail angle and duration, we initially fitted models with interaction terms between head-to-tail angle and duration. For both models, the correlation coefficients of the interaction terms were not significantly different from 0 (head-to-tail angle: *P*=0.069, *N*=33; speed: *P*=0.37, *N*=33), so we eliminated them from the model.

For selected pairs of variables, we performed total-least-squares curve fits with an optimisation method (MATLAB’s `fminsearch`). We normalised both variables to a range of [0, 1]. We fit functions of the form *y* = *c*_1_*x*^*c*_2_^, since we expect a negative power law with an asymptote *y* = 0 when *x* → ∞. For each set of trial coefficients, we calculated the perpendicular distance to the curve for all data points. The squared sum of these distances was used as the objective function of the optimisation, resulting in a set of best-fitting coefficients *c*_1_ and *c*_2_. By bootstrapping with 10,000 repetitions and computing the 2.5^th^ and 97.5^th^ percentile, we calculated 95% confidence intervals of the coefficients.

## Results

### Example of a fast start

We used an automated video-tracking method (Fig. 1) to reconstruct the fast-start motion of a zebrafish larva of 5 days after fertilisation, showing a change in direction of 83 deg, and a maximum speed of 0.15 m s^−1^. The larva curls into a C-shape in stage 1, then unfolds itself in stage 2 followed by a tail beat in opposite direction (Fig. 2A). Over the course of the start, the larva reorients itself from being approximately aligned with the negative *x*-axis of the world reference frame, to swimming in the direction of the positive *y*-axis. In addition, it changes its pitch angle from a nose-down stance to an upward motion.

**Fig. 2.**
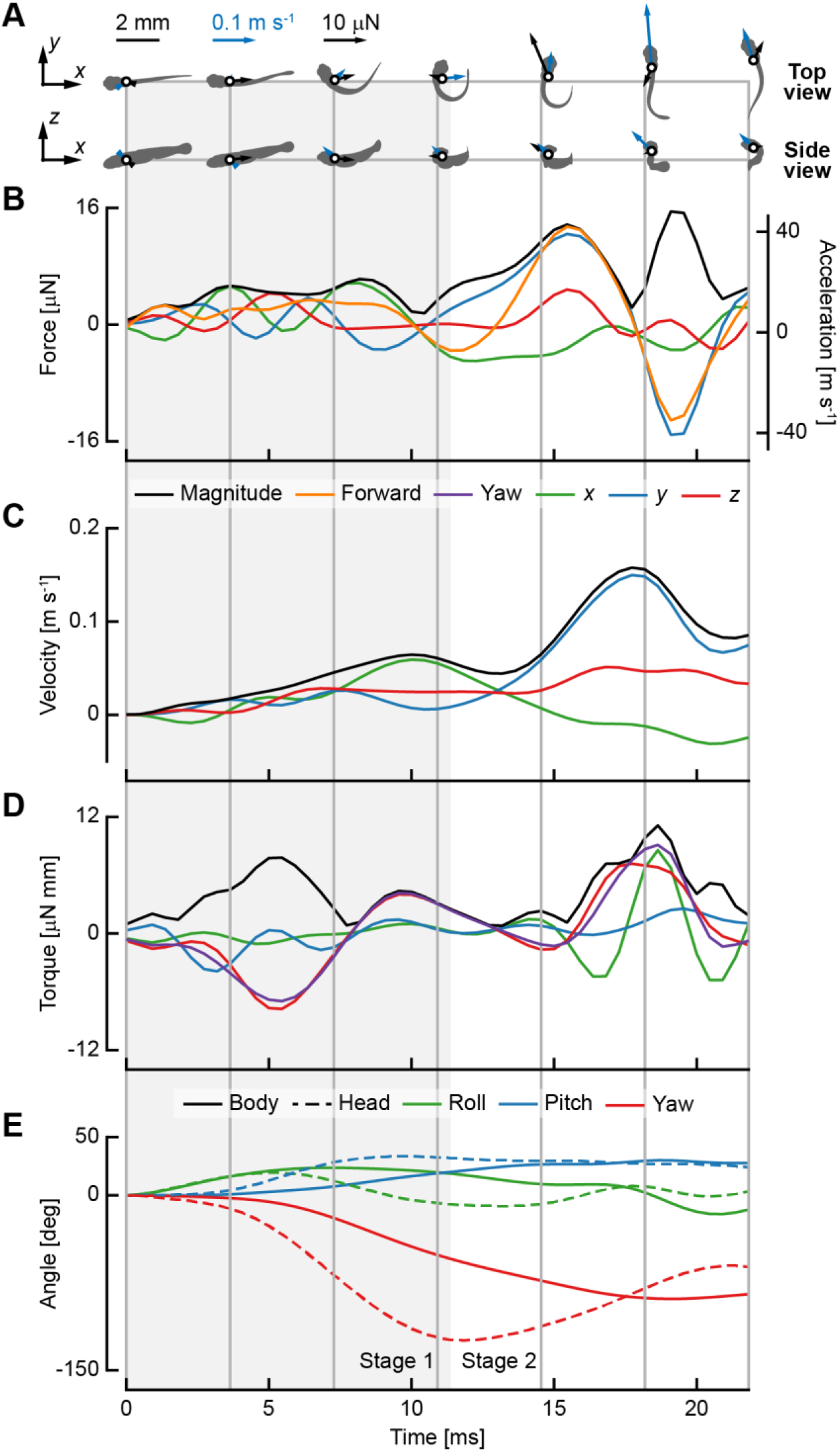
Individual example of a fast-starting zebrafish larvae. Across all sub-panels, the light grey rectangle Indicates stage 1, and the vertical dark grey lines connect the fish shapes in (A) to the time series in (B–E). (A) Projections of the reconstructed fish model in the *x-y* (top) and *x-z* (bottom) plane. The white dot indicates the centre of mass, the blue arrow indicates the instantaneous velocity, and the black arrow represents the instantaneous resultant force. (B) The instantaneous resultant force in *x* (green), *y* (blue), *z* (red), and forward (orange) direction, and the force magnitude (black). The forward direction is defined as the vector pointing in the direction of the instantaneous velocity of the centre of mass. (C) The velocity of the centre of mass in *x* (green), *y* (blue, and *z* (red) direction. (D) The instantaneous resultant torque in *x* (green), *y* (blue), *z* (red), and yaw (purple) direction, and the torque magnitude (black). The yaw torque is defined as perpendicular to the deformation plane of the centre line. (E) The body (solid) angle and head (dashed) Tait-Bryan angles, roll (green), pitch (blue), and yaw (red).

The reconstructed forces vary around 0 in stage 1 of the start, in *x*-, *y*-, and *z*-direction (Fig. 2B). Around the halfway point of stage 2 the force peaks, mainly in “forward” direction (i.e. the direction of the instantaneous velocity vector) – the larva pushes off and produces the largest acceleration resulting in a velocity peak approximately 2 ms later (Fig. 2C). At the same time as the forward peak, an upward (i.e. positive *z*-direction) force peak also occurs, causing an upward velocity of the centre of mass (Fig. 2C). This is followed by a force peak in opposite direction to the velocity, thus decelerating the larva.

The resultant *x*- and *y*-torques are limited in stage 1, but the *z*-torque is considerable (Fig. 2D). The yaw torque is similar to the *z*-torque since the deformation plane is approximately aligned with the *x-y* plane for most of the motion. The first peak of the yaw torque in stage 1 reorients the fish, and is produced while the fish is bending into a C-shape. Later in stage 1, a counter-torque is produced that brakes the reorientation. In stage 2, a higher peak in the same direction as the counter-torque is produced to reorient the fish in the opposite direction during the push-off tail beat.

We determined body angles (Fig. 2E) by integrating the angular velocity that we calculated from the angular momentum and instantaneous moment of inertia. The head angles are defined with the orientation of the stiff head region of the fish. The roll and pitch angle show different dynamics for the head than the body, because the coordinate systems are not aligned so their relative contribution to the out-of-plane orientation changes. The yaw angle is different due to the deformation of the larva – the head angle is not a good indicator for the orientation of the whole larva. The head angle shows large-amplitude variation across the start, while the body angle changes close to monotonously throughout the start, in the direction of reorientation.

### Reorientation and speed

We determined the turn angle of the start by calculating the angle between the initial orientation of the larva and the heading at the end of stage 2. The initial orientation was defined as the unit vector pointing from tail tip to snout, while the heading was defined as the direction of the velocity vector of the centre of mass at the end of stage 2. The “final speed” of the start is defined as the speed of the centre of mass at the end of stage 2. We show the turn angle (Fig. 3A,B) and final speed (Fig. 3C,D) as a function of the head-to-tail angle and start duration. The head-to-tail angle is defined as the angle between the head and the tail at the transition point from stage 1 to stage 2, an indication of the whole-body curvature at the most-curved point. The start duration is computed as the time interval between start initiation and the end of stage 2.

**Fig. 3.**
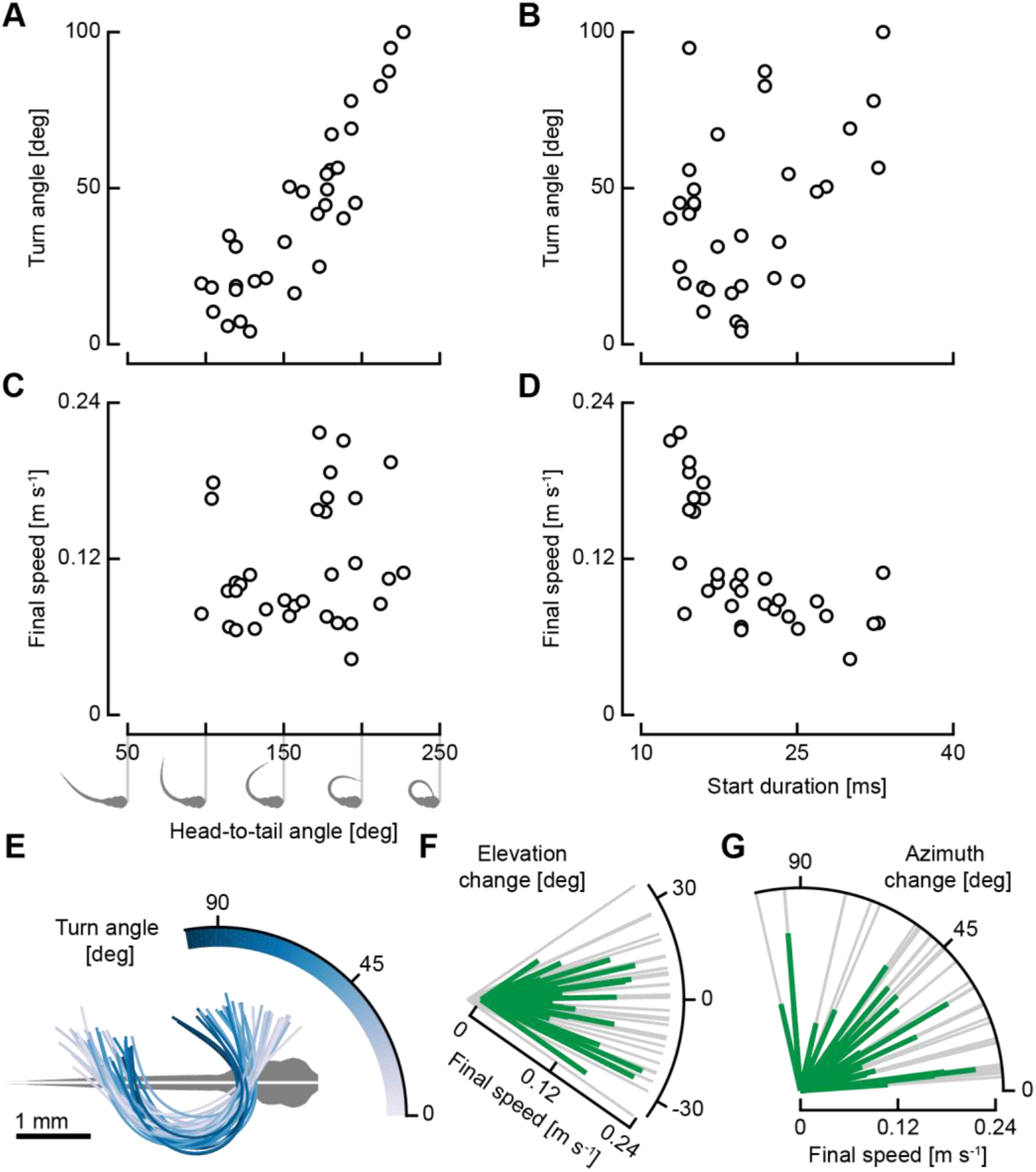
Higher start curvatures increase turn angle, while shorter start durations increase final speed. (A,C) The turn angle (A) and the final speed at the end of stage 2 (C) as a function of the maximum head-to-tail angle, an indication for the total amount of curvature of the body as illustrated below the horizontal axis of C. (B, D) The turn angle (B) and the final speed at the end of stage 2 (D) as a function of the start duration (computed as the time interval between start initiation and the end of stage 2). (E) The shape of the centreline at the transition from stage 1 to stage 2, coloured by the turn angle of the start. (F) The elevation change (curved axis) and final speed at the end of stage 2 (green radial lines) for all analysed starts. (G) The azimuth change (curved axis) and final speed at the end of stage 2 (green radial lines) for all analysed starts.

More strongly curved starts show a higher turn angle – the turn angle is strongly correlated to the head-to-tail angle, with a correlation coefficient of 0.83 (*P*<0.001, *N*=33; bootstrapped 95% confidence interval (CI95%): [0.71, 0.92]). The slope of the correlation is 0.59 (CI_95%_: [0.50, 0.65]) deg of turn angle per deg of head-to-tail angle. In contrast, the turn angle is weakly correlated with the start duration, with a correlation coefficient of 0.19 (*P*=0.032, *N*=33; CI_95%_: [0.028, 0.36]). A longer start duration tends to result in a slightly larger turn angle, at a rate of 0.84 (CI_95%_: [0.125, 1.61]) deg ms^−1^.

Shorter starts have a higher final speed – the final speed is strongly negatively correlated with the duration of the start, with a correlation coefficient of −0.77 (*P*<0.001, *N*=33; CI_95%_: [-0.89, −0.63]). The slope of the correlation is −0.0061 (m s^−1^) ms^−1^ – every millisecond shorter duration will result in a speed increase of 0.0061 m s^−1^. We also fitted a power law to the final speed as a function of start duration, resulting in an exponent of −1.42 (CI_95%_: [-1.87, −1.05]). The final speed is shows a weaker correlation with the head-to-tail angle, with a correlation coefficient of 0.38 (*P*=0.0033, *N*=33; CI_95%_: [0.11, 0.64]). The slope is 4.88 · 10^−4^ (m s^−1^) deg^−1^ (CI_95%_: [1.33 · 10^−4^, 0.82 · 10^−4^]); an increase in head-to-tail angle of 90 deg would result in an increase in final speed of 0.044 m s^−1^.

The centrelines of the fish at the transition from stage 1 to stage 2 are shown in Fig. 3E, transformed to the coordinate system attached to the head of the fish in its initial orientation. The larvae curl up while the centre of mass remains in approximately the same position. The more strongly curved motions show a larger reorientation of the head, as well as a larger turn angle. In general, the head angle at the end of stage 2 is larger than the turn angle at the end of stage 2: the head turns further than the final heading at the end of stage 1, and then turns back over the course of stage 2.

We can divide the total angle change of the body during the start in an elevation angle change (vertical reorientation) and an azimuth angle change (horizontal reorientation), see Fig. 3F,G. The elevation change ranges from −35.0 deg to 34.2 deg (Fig. 3F); the azimuth change ranges from 3.9 deg to 102.7 deg (Fig. 3G). There is no significant correlation between the final speed and the azimuth change (*P*=0.77, *N*=33) or final speed and the elevation change (*P*=0.13, *N*=33).

### Stages of the fast start

We divided the fast start in stages with the same method as Fleuren *et al*. (2018), and analysed the first two stages. The durations of stage 1 and stage 2 are significantly correlated (*P*<0.001, *N*=33; correlation coefficient 0.79, CI_95%_: [0.70, 0.89]). Stage 1 takes on average 52±4.6% of the start until the end of stage 2 (Fig. 4A) – slightly over half of the first two stages is spent bending into a C-shape. No starts were recorded where stage 1 took less than 42% or more than 64% of the start duration. The larvae show a displacement between 3.3–21.1× larger in stage 2 compared to stage 1 (Fig. 4B). Also the speed is larger, both total speed (Fig. 4C; 1.5–5.6×) and speed in the direction of the final heading (Fig. 4C, “forward”; 1.7–23.6×). The total speed in stage 1 is higher than the “forward” speed – the centre of mass moves slightly in stage 1, but not much in “forward” direction (i.e. in the direction of the velocity at the end of stage 2).

**Fig. 4.**
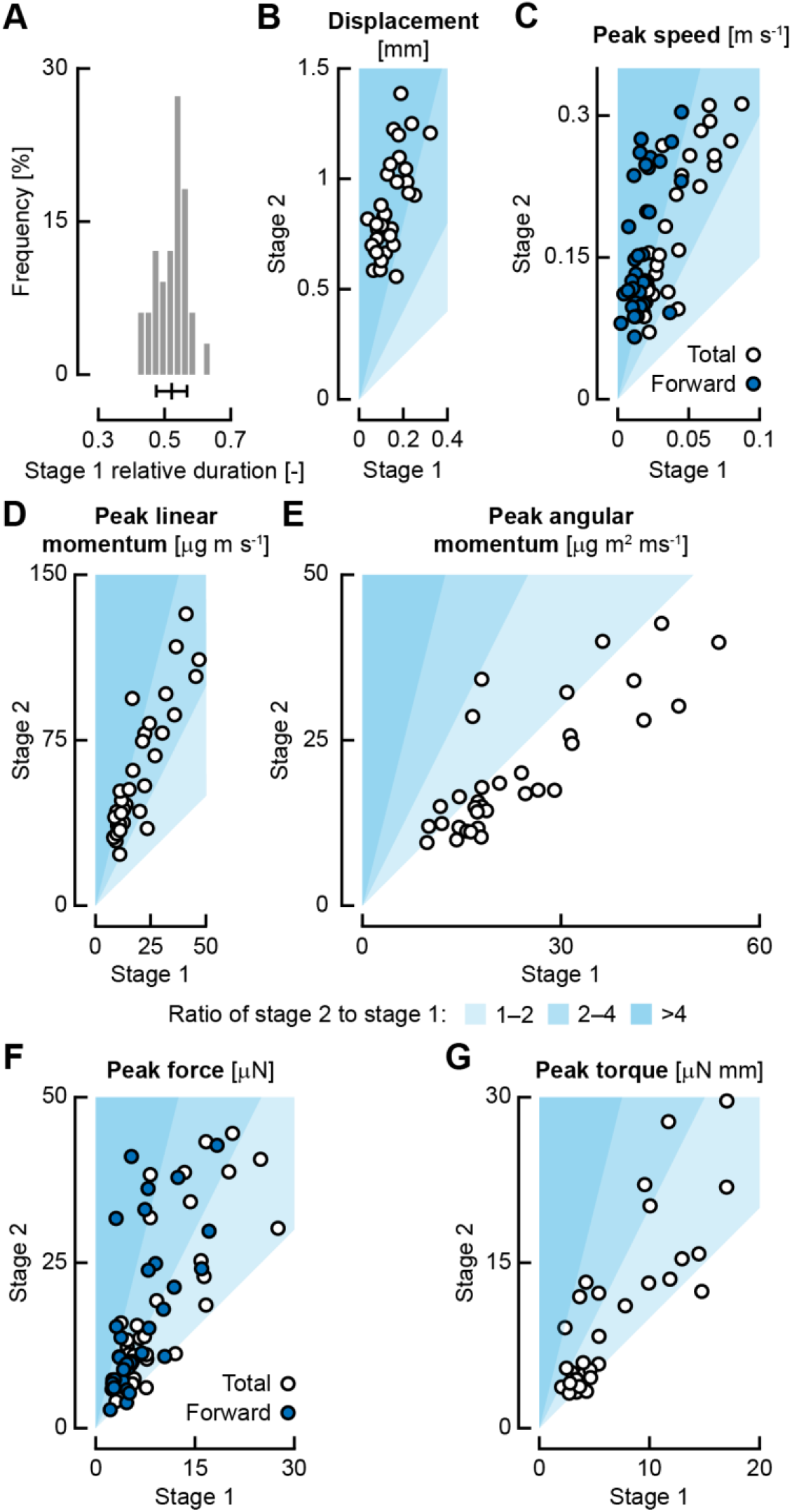
Stages of the fast start: highest speeds, forces, and torques occur in stage 2. (A) The duration of stage 1 with respect to the duration of the start (computed as the time interval between start initiation and the end of stage 2), the histogram indicates the frequency of each bin as a percentage of all starts. The bar below the histogram shows the mean and one standard deviation. (B–G) The white region indicates where the values in stage 2 are lower than stage 1, in the light blue region, the values for stage 2 are 1–2× higher, in the medium blue region 2–4×, and in the dark blue region >4×. The contribution of stage 1 is on the horizontal axis, the contribution of stage 2 on the vertical axis. (B) Net displacement, i.e. the reduction in distance to the final position of the centre of mass at the end of stage 2. (C) Peak speed; white dots indicate the total speed, blue dots indicate the speed in the direction of the final heading. (D) Peak linear momentum. (E) Peak angular momentum. (F) Peak force; white dots indicate the total force, blue dots indicate the force in the direction of the final heading. (G) Peak torque.

In all cases, the peak linear momentum is larger in stage 2 than in stage 1 (Fig. 4D; 1.5–5.6×), while the peak angular momentum is often smaller in stage 2 than in stage 1 (Fig. 4E; 0.58– 1.9×). Stage 1 therefore often shows higher angular velocities than stage 2. In most cases, the peak force is higher in stage 2 than in stage 1 (Fig. 4F), this holds for both the total force (0.80–4.6×) and the “forward” force (i.e. in the direction of the velocity at the end of stage 2; 0.82–10.3×). Not much force is produced in stage 1, especially in the direction of the start – the acceleration is mostly visible as an undirected wiggling of the centre of mass. In most cases, the torque is also higher in stage 2 than in stage 1 (Fig. 4G; 0.78–3.9×), but the ratio is smaller than that of the speed and forces; some sequences even show higher torques in stage 1 than stage 2. The higher torques in stage 2 are presumably produced by the higher forces during the push-off.

### Reorientation

Stage 1 has a significantly higher contribution to the yaw angle change than stage 2 (Fig. 5A; *t*-test, *P*<0.001, *N*=33); on average the contribution of stage 1 is 28.7±13.7 deg higher than the contribution of stage 2. For smaller total yaw changes, stage 2 might have a negative contribution, undoing part of the reorientation of stage 1. Phase plots of the yaw angle (Fig. 5B) show that starts with relatively small turn angles generally have a negative contribution of stage 2 to the body yaw angle, while for large turn angles the body yaw angle changes almost monotonously. In contrast, the head yaw angle shows considerably larger variation over the fast start than the body angle, reaching a maximum near the end of stage one, before rotating in opposite direction in stage 2.

For all fast starts, we averaged the linear momentum, angular momentum, and change in moment of inertia normalised by their maximum value (Fig. 5C–E). The linear momentum (Fig. 5C) reaches a small peak in stage 1, followed by a much larger peak in stage 2, where peak speed is reached. In contrast, the angular momentum (Fig. 5D) shows its largest peak in stage 1, followed by a lower peak in stage 2. The large peak in angular momentum just proceeds to the dip in moment of inertia (Fig. 5E). A combination of large angular momentum and low moment of inertia leads to a high angular velocity, indicating a strong reorientation in stage 1. After the peak, the angular momentum reduces, indicating that the yaw rotation is braked by a counter-torque before rising again as the fish beats its tail in the opposite direction.

**Fig. 5.**
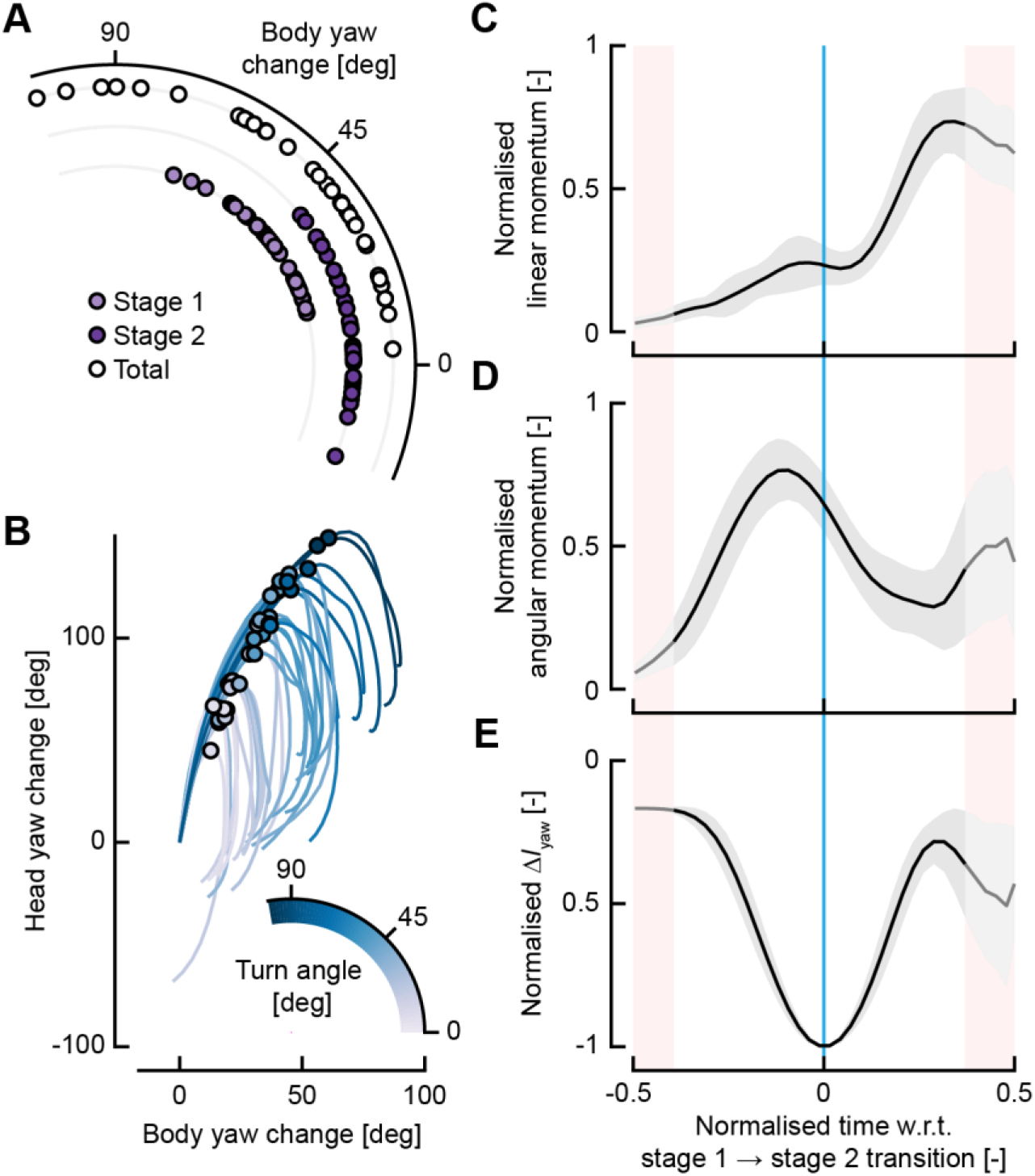
Reorientation during the fast start. (A) Change in yaw angle in stage 1 (light purple), stage 2 (dark purple), and the complete start (white). (B) Phase plots of the change in head yaw angle to the change in body yaw angle, coloured by the turn angle of the start (the angle between the initial orientation of the fish and the direction of the velocity vector at the end of stage 2). The dots indicate the transition point from stage 1 to stage 2. (C–E) The black line indicates the mean, with ± 1 standard deviation shown by the grey area. The red bands at the left and right indicate time points for which not all data is present due to differences in the relative length of stage 1 and 2. The blue line indicates the transition point from stage 1 to stage 2. (C) The linear momentum profile over the start, normalised by its maximum value and aligned to the transition from stage 1 to stage 2 before averaging over all starts. (D) The angular momentum profile over the start, normalised by its maximum value and aligned to the transition from stage 1 to stage 2 before averaging over all starts. (E) Changes in moment of inertia in the deformation plane (A*I*_yaw_), normalised by its minimum value and aligned to the transition from stage 1 to stage 2 before averaging over all starts.

### Propulsion in stage 2

We calculated the speed of the tail as the speed averaged over the posterior 10% of the body, relative to the speed of the centre of mass. The peak tail speed over the fast starts tends to increase with decreasing duration of the motion (Fig. 6A), with a correlation coefficient of - 0.67 (*P*<0.001, *N*=33; CI_95%_: [-0.79, −0.54]). For every millisecond of decrease in duration, the peak tail speed increases by 20.3 m s^−1^ (CI_95%_: [16.3, 23.9]). In addition, we fitted a power law to the tail speed as a function of start duration, resulting in an exponent of −1.27 (CI_95%_: [-1.68, - 0.96]). Since much of the propulsive force is produced at the tail, which moves in opposite direction to the velocity of the centre of mass (Fig. 1A), the peak force tends to increase with increasing peak tail speed (Fig. 6B), with a correlation coefficient of 0.85 (*P*<0.001, *N*=33; CI_95%_: [0.70, 0.94]). The slope of the correlation is 58.4 μN (m s^−1^)^−1^ (CI_95%_: [48.3, 64.7]). In this way, a decrease in duration leads to an increase in tail speed, and hence a corresponding increase in propulsive force, and therefore leads to an increase in escape acceleration.

**Fig. 6.**
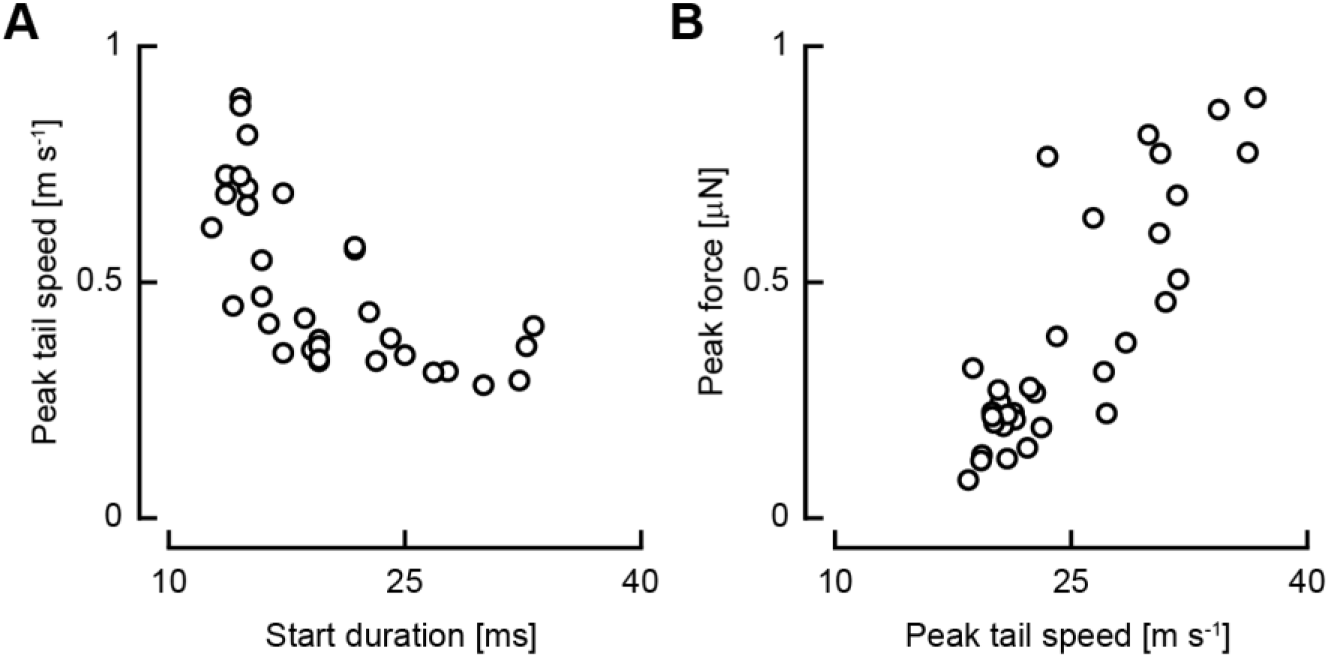
Higher tail speeds with shorter durations lead to higher forces. (A) The peak tail speed in stage 2 as a function of the start duration. (B) The peak force in stage 2 as a function of the peak tail speed in stage 2.

### Pectoral fin use during the fast start

For each time point in the fast start, we manually indicated whether or not the pectoral fins were abducted. During high-speed starts, the pectoral fins remain adducted for the entire duration of the start, while during slower starts, they are abducted for part of the start (Fig. 7A). Whether the pectoral fins are abducted during a start does not depend on the turn angle (Fig. 7A). In starts where the fins were used, they were first abducted in stage 1 after 8.2±6.0% of the start duration (Fig. 7B). They were then adducted in stage two, after 75±8.3% of the start duration, resulting in an average duration of pectoral fin abduction of 67±8.6% of the start.

**Fig. 7.**
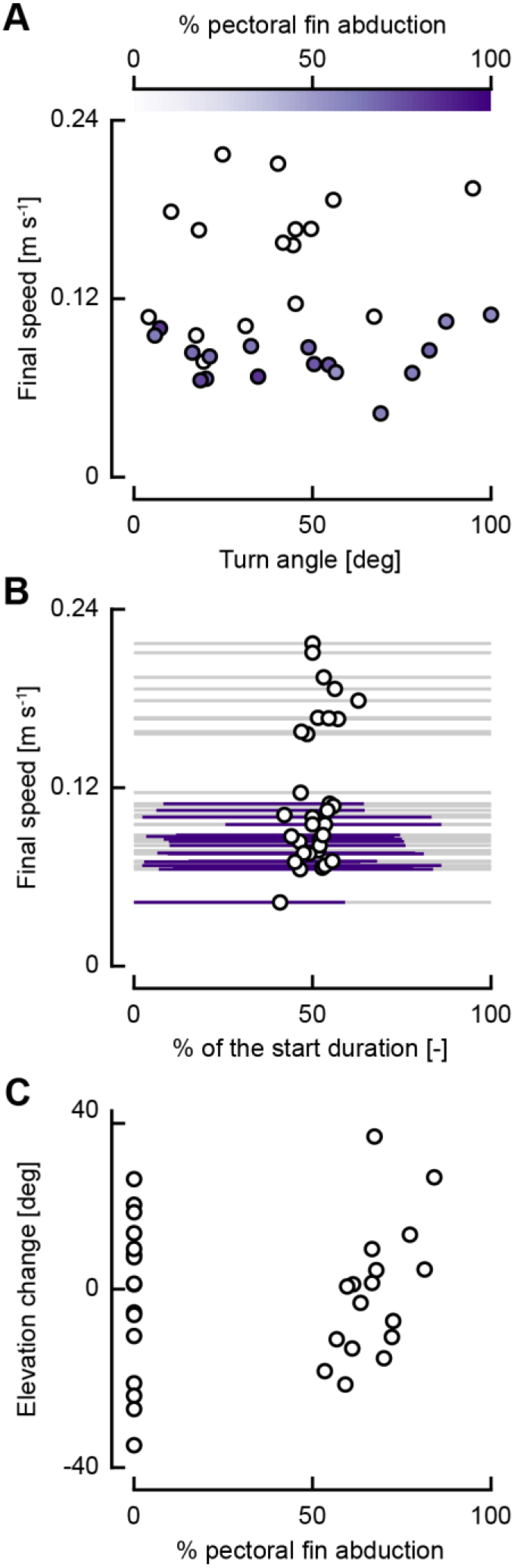
Pectoral fins are only abducted during low-speed starts and help produce elevation change. (A) The percentage of the start duration (computed as the time interval between start initiation and the end of stage 2) that the pectoral fins are abducted (colours) in the turn angle – final speed landscape. (B) The relative time as a fraction of the start that the pectoral fins are abducted (purple lines) as a function of the speed. Each grey line indicates a start, with the transition point between stage 1 and 2 indicated by the dot. (C) The elevation change as a function of the percentage of the start that the pectoral fins are abducted.

In starts where the fins are used, the fraction of the start that they are abducted correlates significantly with the change in elevation (*P*=0.0252, *N*=17), with a correlation coefficient of 0.54 (CI_95%_: [0.15, 0.83]). In starts where the fins are not used, large elevation changes could also be produced – the mean elevation change between starts with and without fins are not significantly different (two-sample *t*-test, *P*=0.82, *N_1_*=17, *N_2_*=16).

## Discussion

We reconstructed the three-dimensional motion of zebrafish larvae of 5 days after fertilisation during C-start escape responses and reconstructed linear and angular momentum, and forces and torques. We consider the results of the analysis in the context of the functional demands of the start, as outlined in the introduction.

### Producing acceleration

The primary demand of a fast start is to accelerate the body, both linearly and rotationally. This acceleration is produced by a large force peak in stage 2 (Fig. 2B, Fig. 4F), causing an increase in linear momentum, and hence speed (Fig. 2C, Fig. 4D, Fig. 5C). Although the body is prepared for the propulsive stroke by curling up in stage 1, the body curvature (as expressed with the head-to-tail angle) correlates with the speed relatively weakly (Fig. 3C). In contrast, the speed shows a strong inverse correlation with the duration of the start, with a power law exponent of −1.42: shorter starts lead to higher speeds (Fig. 3D). The durations of the stage 1 and stage 2 do not vary independently (Fig. 4A). Hence, shorter start durations lead to shorter durations of stage 2, resulting in an increase in tail speed (Fig. 6A) with a power law exponent of −1.27, and a resulting increase in force (Fig. 6B).

To produce these forces, fish produce fluid-dynamic jets. During stage 1, fish larvae produce a jet flow into the C-shape (Li et al., 2014; Müller et al., 2008). A CFD simulation of a single zebrafish larva swimming sequence (Li et al., 2012) showed that initially this mainly produces a torque that reorients the fish. The jet is then reoriented along the body in stage 2, where it produces propulsive force, in agreement with our reconstructed resultant forces (Fig. 2B). Adult bluegill sunfish show a similar flow pattern in velocity field measurements (Tytell and Lauder, 2008).

Based on numerical simulations it has been found that the motion of the larval C-start was near-optimal for maximising escape distance in a given time (Gazzola et al., 2012) – a measure that corresponds to maximising the mean acceleration during a start from a standstill. They also found that a higher curvature could result in a higher escape distance, given a start duration; this corresponds to the weak correlation that we find for speed with head-to-tail angle. For (near-)cyclic swimming of larval fish, the swimming speed was found to increase with increasing tail-beat frequency and to a lesser extent amplitude (Van Leeuwen et al., 2015). The fast start duration is the equivalent of the frequency, while the head-to-tail angle is connected to the tail-beat amplitude. Hence, we see similar effects on the speed in cyclic swimming as in fast starts.

### Reorienting the body

The larvae produce a wide range of escape directions (Fig. 3F,G), both in azimuth and, to a lesser extent, in elevation. The turn angle of a start correlates strongly with the head-to-tail angle: more strongly curved starts tend to show a larger turn angle (Fig. 3A,E). The turn angle correlates weakly with the duration of the start (Fig. 3B), where longer starts show a slightly larger turn angle. Hence, large turn angles do not take much more time to produce than small turn angles. In adult fish, the start duration correlates more strongly to the escape angle (angelfish: Domenici and Blake, 1991; goldfish: Eaton et al., 1988). This suggests a difference in reorientation between adults and zebrafish larvae: adults seem to use an approximately fixed turn rate, while larval zebrafish increase turn rates with increasing turn angles.

The changes in escape angle are mostly produced in stage 1 (Fig. 5A), despite lower peak torques (Fig. 4G). The yaw torque is consistently in the direction of turning during the first part of stage 1 (Fig. 2D), causing the angular momentum to show its largest peak in stage 1 (Fig. 5D), while the moment of inertia is close to its minimum (Fig. 5E). The high angular momentum combined with a low moment of inertia leads to a high angular speed, allowing large turn angles. At the end of stage 1, the torque reverses sign (Fig. 2D), thus reducing the angular momentum. Together with the increase in moment of inertia (Fig. 5E), this decreases the angular speed. The torque then decreases until the end of stage 2, where the torque increases again, rotating the fish in opposite direction (Fig. 2D). This reorienting torque and the following counter-torque were shown to be caused mainly by pressure forces, while the largest shear forces were found at the head, and counteracted the initial reorienting torque (Li et al., 2012)

Previous studies of adult fish have shown that the turn angle of the head in stage 1 correlates with the turn angle during the complete fast start (Domenici and Blake, 1993; Eaton et al., 1988; Fleuren et al., 2018). This has also been found for fast starts of zebrafish larvae (Nair et al., 2015). Danos and Lauder (2007) analysed routine turns of zebrafish larvae, for which they created a model where only the body bending caused a change in head angle, resulting in a large underprediction of the escape angle. They suggested that the additional effect is caused by fins. In fast starts, however, the pectoral fins cannot explain the reorientation torque as they are adducted at high speeds, even for large turn angles (Fig. 7A). Without fins, fish have been shown to produce a yaw torque in the first stage of the start (Li et al., 2012; Song et al., 2018). This torque is mainly produced by pressure forces at the tail, which has a much larger lever arm with respect to the centre of mass than the pectoral fins.

### Alternatives to the head angle

The head angle change after stage 1 is connected to the head-to-tail angle of the fish due to the stereotypical nature of the C-bend (Fig. 3E). The tail excursion of zebrafish larvae was found to correlate with the head yaw angle (Nair et al., 2015), so the head-to-tail angle correlates with the head yaw angle. Rather than use the head angle to indirectly indicate the curvature of the start, we use the head-to-tail angle as a more direct indicator of the whole-body curvature. Since the posterior part of the fish produces much of the reorienting torque (Li et al., 2012; Song et al., 2018), it is useful to consider the complete body when analysing at escape direction.

Furthermore, rather than using the head angle as an indicator for orientation (Domenici and Blake, 1993; Eaton and Emberly, 1991; Nair et al., 2015), we use the “body angle”, that we calculate from the mass distribution. The head angle is not representative of the heading of the fish: they differ considerably across most of the fast start (Fig. 2E, Fig. 5B). The body angle is more difficult to quantify than the head angle, as it requires a three-dimensional mass distribution model of the fish, and reconstructed kinematics of high accuracy (Van Leeuwen et al., 2015). Nonetheless, it is worth calculating when analysing reorientations, as it gives a much more accurate representation of the reorientation of the fish mass. In the absence of body angles, the head angle cannot be used to replace it, as it shows completely different dynamics.

### Control of the fast start

The turn angle and final speed seem to be adjusted mostly independently for C-starts of zebrafish larvae. The turn angle can be adjusted with the head-to-tail angle (i.e. body curvature), having relatively limited effect on the escape speed (Fig. 3A,C). The escape speed can be adjusted with the start duration, having a limited effect on the escape angle (Fig. 3B,D). In adult goldfish, the escape trajectory was found to be controlled by the relative size of the initial and second contractions and the timing between them with minimal feedback from sensors (Foreman and Eaton, 1993). Assuming that starts are controlled similarly in larval zebrafish, the head-to-tail angle and start duration are presumably a direct result of these parameters, and might be used as proxies for them.

The duration of stage 1 and stage 2 vary concomitantly (Fig. 4A), also previously found for two species of adult fish (Webb, 1975) and zebrafish larvae (Nair et al., 2015). The larvae do not individually tune the duration of stage 1 and stage 2 to adjust the angle and speed of their escape. At a given escape speed, smaller head-to-tail angles are produced by turning more slowly, rather than turning at the same rate but shorter. Furthermore, the duration of stage 2 is not shortened independently of stage 1 to increase the tail speed, and hence propulsive force. This might suggest a limitation on how quickly the tail-beat duration can be changed from one tail-beat to the next.

The elevation of the start has been found to be controlled by dorsoventral excursions of the midline (Nair et al., 2015). In the slow starts where the pectoral fins were used, the amount of time that the pectoral fins were abducted correlates with the elevation change (Fig. 7C). Larvae of 5 dpf naturally show a nose-down pitch moment (Ehrlich and Schoppik, 2017), so a hydrodynamic torque must be produced to counteract this for positive, or perhaps even less-negative elevation changes. The action of the pectoral fins is an additional effect to the dorsoventral tail excursion, since starts without pectoral fin abduction do not produce significantly different elevation changes. The pectoral fins are only used during relatively slow C-starts (Fig. 7A). At lower speeds, perhaps the required pitch torques cannot be produced by the body alone, requiring help of the pectoral fins. In contrast, at high speeds, the body is able to produce sufficient pitch torque, and can adduct the fins to reduce drag to achieve a higher escape speed.

### Timing the start

The importance of fine-tuning the speed and direction of the escape depends on speed of the predator relative to the prey. When the speed of the predator is close to the speed of the prey, faster starts will result in greater survival probability (Walker et al., 2005). However, for a much faster or much slower predator than prey, the speed and direction are less important than for intermediate predator speeds (Soto et al., 2015). Zebrafish larvae have been stated to be mostly in the “slow predator” regime, where escape timing is the dominant parameter (Stewart et al., 2013) influencing escape performance, although below strong reductions (>50%) in escape speed, the probability of escape from the predator’s suction flow drops rapidly (Nair et al., 2017).

Zebrafish larvae show a relatively long stage 1 (Fig. 4A), in which hardly any propulsion is produced (Fig. 4B,C,F), reducing the mean acceleration of the start. However, if the zebrafish detects the threat sufficiently early, it can initiate stage 1 of the fast start to begin stage 2 at the optimal moment. Hence, the relatively long duration of stage 1 without significant propulsion might not be a disadvantage in escaping predators for zebrafish larvae.

### Contributions of stage 1 and stage 2

Stage 1 is necessary to prepare the body for acceleration, but takes up, on average, over half the time of a fast start without providing much propulsion (Fig. 4). The displacement is much larger in stage 2 (Fig. 4B), as is the peak speed, especially in the direction of the final heading (Fig. 4C). Stage 2 shows a larger linear momentum than stage 1 (Fig. 4D), as well as a larger peak force (Fig. 4F). In contrast, the angular momentum is often smaller in stage 2 than in stage 1 (Fig. 4E), despite the generally higher peak torques in stage 2 (Fig. 4G). The torques are more consistently in the direction of reorientation in stage 1, allowing the angular momentum to build to a higher value.

The role of stage 1 and stage 2 in the fast start has been the subject of on-going debate. The first stage has often been called purely preparatory (Domenici and Blake, 1997; Hertel, 1966; Weihs, 1973). Stage 1 prepares the body for stage 2: its preparatory role is clear (Fleuren et al., 2018). In addition to the preparatory function, it has also been argued that stage 1 may contribute significantly to propulsion (Fleuren et al., 2018; Tytell and Lauder, 2008; Wakeling, 2006). For bluegill sunfish, 37.2±0.6% of linear momentum is produced after stage 1 (Tytell and Lauder, 2008); for the larval zebrafish this is somewhat lower at 27.8±8.2% (Fig. 5C). Based on the linear momentum, there is some propulsion component in stage 1, but the displacement, speed, peak linear momentum, and peak forces are all considerably lower compared to stage 2 (Fig. 4). Arguably, the preparatory role of the start, including reorientation, is more important for zebrafish larvae than the propulsive role.

## Conclusions

In this article, we analysed the dynamics of the fast start of zebrafish larvae at five days post fertilization. We confirm that early-development larvae can produce effective escape response in a wide range of directions (both azimuth and elevation) and speeds. The larvae seem to be able to adjust the direction and speed of their escape close to independently. They adjust the escape angle mostly with the extent of body curvature, while the escape speed is adjusted mostly with the duration of the start. Apart from its preparatory role, stage 1 is used to produce most of the reorientation, while stage 2 produces most of the acceleration of the centre of mass. This shows that despite their early stage of development, zebrafish larvae meet the functional demands for producing effective escape responses.

## Acknowledgements

We thank the staff of the Carus fish facilities for providing the zebrafish larvae. We thank Kas Koenraads for his assistance during the experiments.

## Data availability

Data will be made available in a public repository.

